# Glucocorticoids mediate egg rejection in a brood parasite host

**DOI:** 10.1101/818864

**Authors:** Mikus Abolins-Abols, Mark E. Hauber

**Affiliations:** Department of Evolution, Ecology, and Behavior, School of Integrative Biology, University of Illinois, Urbana-Champaign, IL 61801, USA

**Keywords:** avian brood parasitism, egg rejection, corticosterone

## Abstract

Avian brood parasites and their hosts are engaged in a coevolutionary battle that can result in the evolution of sophisticated trickery by parasites and novel defence behaviours in hosts. Despite the clear evolutionary and ecological significance of host behaviour, however, we know very little about the mechanisms that regulate host defences, which limits our understanding of both inter- and intraspecific variation in host responses to parasitism. Here we tested whether corticosterone, a hormone known to be upregulated in hosts exposed to parasitism, also mediates one of the most frequent host defences – the rejection of foreign eggs. We experimentally reduced corticosterone levels in free-living brood parasite hosts, American robins *Turdus migratorius*, using mitotane and found that the likelihood of model egg rejection was significantly lower in the mitotane-treated birds relative to the sham-treated birds. These results demonstrate a causal link between glucocorticoids and egg rejection in hosts of avian brood parasites, but the physiological and sensory-cognitive pathways that regulate this effect remain unknown.

## 1. Introduction

In obligate avian brood parasitism, parasites exploit host parental care by forcing the foster parents to provision unrelated offspring [1,2]. To mitigate the negative fitness consequences of parasitism, hosts can evolve resistance strategies, such as the ability to recognize and reject foreign eggs or young in the nest [3,4], or tolerance strategies, such as the ability to withstand the physiological costs of caring for, or coexisting with, a parasitic nestling [5]. Although much is known about the perceptual cues and behavioural responses that hosts employ to recognize and reject brood parasitic stimuli [6], the physiological mechanisms that underlie host responses to brood parasites remain poorly understood [7].

A handful of recent studies suggest that host responses to avian brood parasitism are, in part, mediated by steroid hormones [8]. For example, host parents caring for a parasitic chick show higher stress-induced corticosterone levels compared to parents caring for only their own young [9]. An increase in the baseline corticosterone levels in hosts can occur directly in response to encountering a foreign egg in the nest [10]. Furthermore, parasitism can alter the endocrine milieu of host offspring, either through an increase in maternally-deposited yolk androgen concentrations [11], or by an increase in the baseline corticosterone levels in host nestlings during competition with parasitic chicks [12]. The physiological and behavioural consequences, and the adaptive value of these hormonal changes, if any, are as of yet unclear, because glucocorticoids in general, and corticosterone in particular, can regulate diverse and even opposing functions [13].

Elevated corticosterone levels in hosts in response to foreign eggs suggests a possible role of this hormone in mediating host resistance or tolerance strategies. Here we tested the hypothesis that egg rejection, a widespread host defence against brood parasites, is mediated by circulating corticosterone. Specifically, we experimentally lowered plasma glucocorticoid levels using mitotane injections in an egg-rejecter host species and asked whether this treatment shifted the probability of non-mimetic foreign egg rejection compared to a sham-treatment. Because corticosterone can mediate vigilance in vertebrates [14] and suppress parental behaviours in birds [15,16], we predicted that inhibition of glucocorticoid synthesis would reduce foreign egg rejection.

## 2. Methods

### (a) Field site and species

We studied wild American robins *Turdus migratorius*, an occasional host to the obligate brood-parasitic brown-headed cowbirds *Molothrus ater*, at Wandell’s Tree Nurserty in Champaign County, IL, USA, during the summer of 2019 (details of the study area are given in [17,18]). We searched for robin nests daily. Prior to the experiment, we monitored the content of focal nests every third day, assuming the clutch was complete when the egg number did not change within 24 hrs [19]. For this study, we focused only on female robins, because in this and other species with female-only incubation they are the only sex responsible for egg rejection [20].

### (b) Treatment validation

Mitotane is a glucocorticoid synthesis inhibitor that has been shown to consistently reduce baseline [21,22] and stress-induced [22–24] corticosterone levels in birds, often to non-detectable levels. We first tested if the effect of mitotane on corticosterone levels in American robins parallels that seen in other species. We caught wild egg-laying or incubating robin (n=8) females and took a baseline blood sample from the brachial vein within 3 min of capture (mean start time = 130 sec; mean end time = 167 sec). Blood was stored on ice and centrifuged at 8000 RPM within 2 hrs to separate plasma. Plasma samples were kept on ice until frozen at –80 °C 4 hrs later. We then injected the pectoral muscle of 4 females with 34 mg mitotane (Sigma-Aldrich, Cat. No. 25925), dissolved in 400 µl sterile peanut oil (Acros Organics, Cat. No. 416855000, dosage 400 mg/kg), following guidelines for the maximum mitotane dosage used in other species [21,22]. Four sham females were injected with a pure peanut oil vehicle (400 µl).

Recapturing wild robins within a day in most cases is not possible, because they become extremely vary of mist nets and humans. We therefore temporarily transferred the injected birds to captivity. We housed birds singly overnight in 40×40×34 cm cages, providing them with ad libitum water, earthworms, bananas, and crushed dry dog food. The following day, we again collected their blood within 3 min of capture (mean start time = 94 sec; mean end time = 120 sec). Captivity was an intense stressor: despite a constant access to food, all birds lost mass, although, on average, they lost less mass in the mitotane (8.1%) than in the sham (14.3%) treatment (two-tailed t-test, n=6, p=0.01).

To test the effect of mitotane on corticosterone synthesis, we analysed plasma corticosterone using an enzyme immunoassay (Cayman Chemical, Cat. No. 501320). Validation details and methods for this assay using robin plasma are published elsewhere [25]. All samples were assayed in duplicate on the same plate, with an intraplate coefficient of variation of 6.7%.

### (c) Hormone manipulation in the wild

We captured incubating robin females (n=61) at their nests using a mist net between 6-10 am, soon after they had completed their clutches (median 2 days, range 0-5 days after clutch completion). We first took a 450 µl blood sample as part of a different investigation. We then injected the bird either with mitotane or sham treatment, as described above. We also took standard morphometric measurements, including age (using wing feather coloration [26]), mass (nearest g), tarsus (nearest 0.1 mm), wing (nearest mm), and pectoral muscle condition [27]. We then fitted females with a USGS band and 3 plastic colour bands (Avinet) to facilitate individual identification in the field, following which the birds were released. An unanticipatedly large number of birds abandoned their nests after the injections (see Results and the Ethics statement), which resulted in a substantial reduction in the final sample size (n=37) compared to the initial captures.

### (d) Experimental parasitism with model eggs

American robins reject the majority of natural cowbird [28] or model cowbird-like eggs [3], whereas they show more variable responses to egg colours near their rejection threshold, which lies between brown-beige and robin-blue colours [29]. Importantly, robins show variable but individually repeatable responses to deep-blue cowbird-sized model eggs (figure 2 inset; for details, see [30]). We therefore used deep-blue 3D-printed eggs to investigate the effect of the injection treatment on egg rejection.

A day after the mitotane or sham injections, we added one deep-blue model egg to the focal nest. We verified the identity of the female using band colours, and checked the temperature of eggs to establish that the nest was active. We did not remove any robin eggs, because *Turdus* thrushes show the same response to model eggs regardless of whether their own eggs are removed [31]. We returned to the nest one day after the addition of the model egg to record whether it was accepted (present in the nest) or rejected (missing). We again checked if the nest was active. If the female was not observed during these visits, we returned to the nest later to confirm female identity. Robins do not abandon their nests following experimental parasitism (relative to control eggs; [32]). Depredated nests were excluded from the analysis.

### (e) Statistical analyses

In the treatment validation study, baseline corticosterone levels were not normally-distributed, therefore we used Mann-Whitney U-tests to assess the differences in hormone levels between mitotane and sham treatments.

We found that life history, seasonal, and morphological variables were not different between the treatment groups (all p>0.05, data not shown). Because the treatments were randomized across individuals and time, we tested the effect of the injections on bird behaviour using Fisher’s exact tests.

## 3. Results

In the treatment validation study, females had similar corticosterone levels prior to the injections (two-tailed Mann-Whitney U test, n=8, U=6, p=0.686). After a day in captivity, as predicted, the mitotane-treated birds (n=4) showed a smaller increase in corticosterone compared to the sham-treated birds (n=4; one-tailed Mann-Whitney U test, U=2, p=0.057, figure 1).

**Figure 1.**
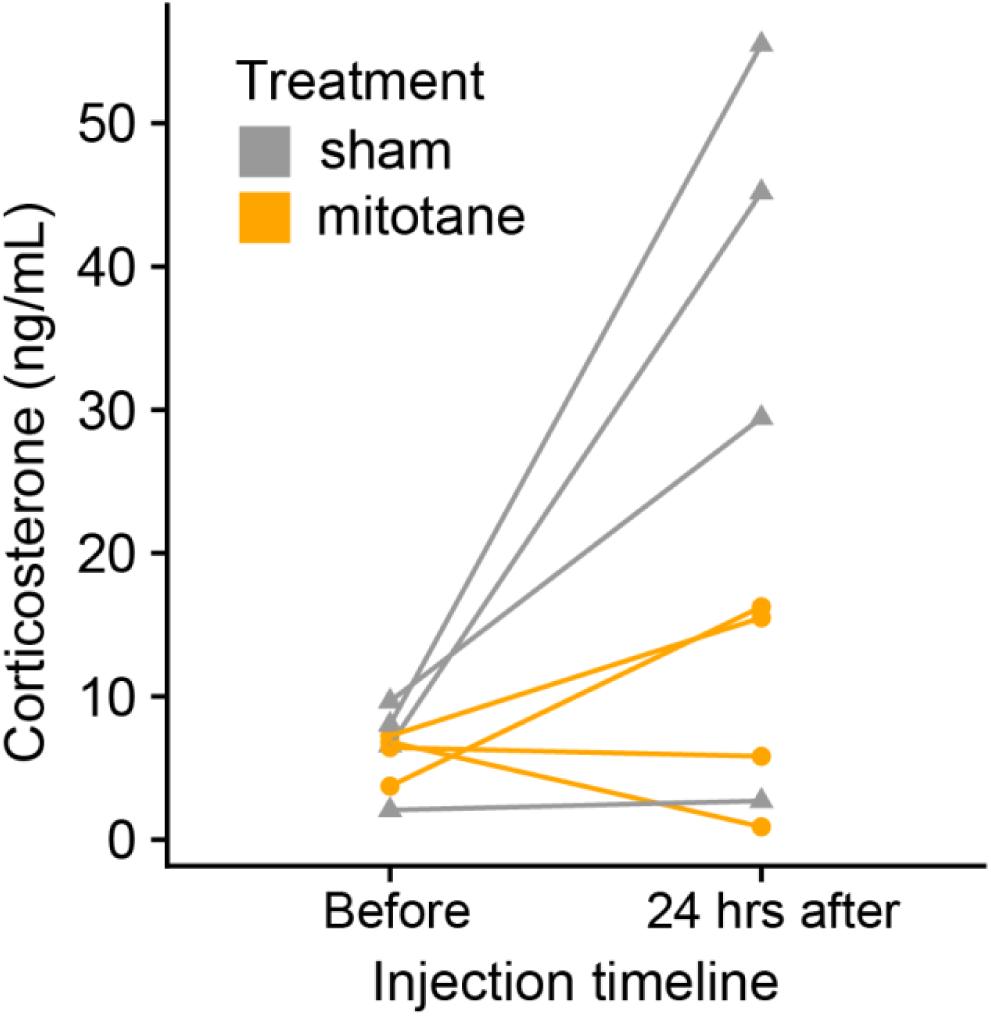
Treatment validation in wild-caught, captive American robin females. Mitotane-treated birds (orange round dots) showed lower increase in corticosterone levels compared to the sham-treated birds (grey triangles). Samples taken from the same individual are connected with lines.

Twenty-four out of 61 females (39%) abandoned their nests within a day of the experimental injections. The probability of nest abandonment did not depend on the treatments (two-tailed Fisher’s Exact test, p=0.601).

The mitotane-treated birds were significantly more likely to accept the model egg (15 out of 20, 75%) compared to the sham-treated individuals (7 out of 17, 41.2%; two-tailed Fisher’s Exact test, n=37, p=0.050, figure 2).

**Figure 2:**
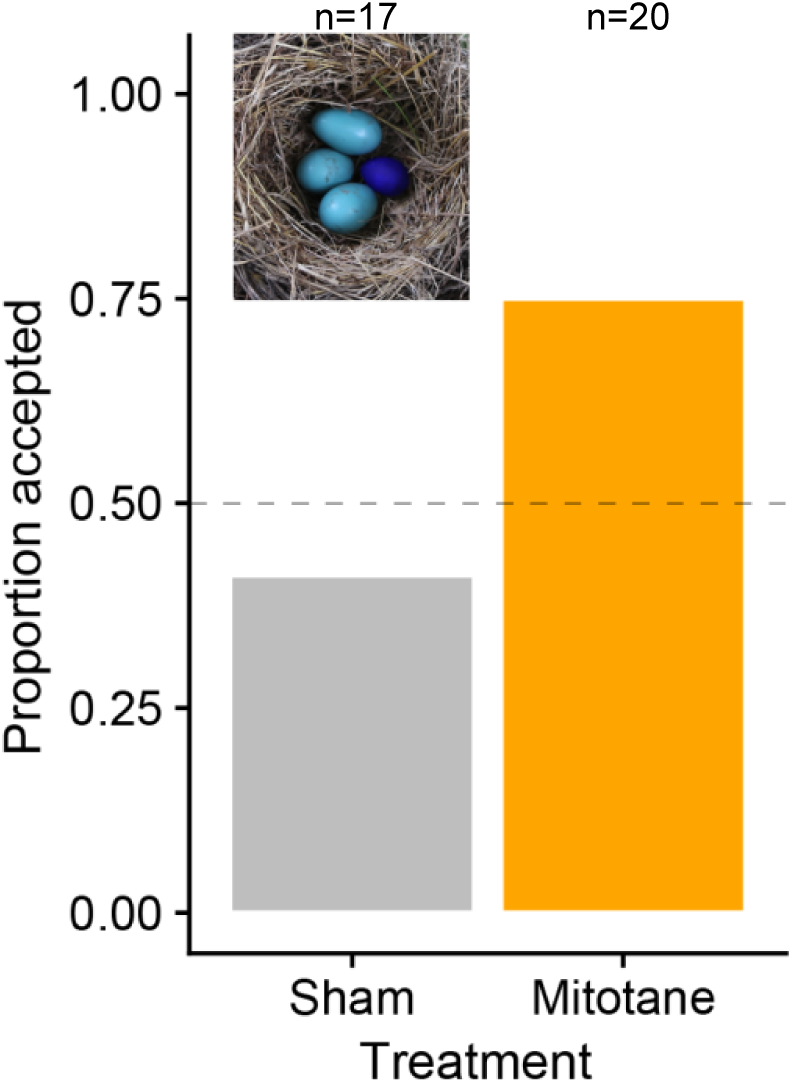
The effect of mitotane on non-mimetic model egg acceptance. Mitotane-treated birds (orange) were significantly more likely to accept model eggs compared to the sham-treated birds (grey). The dotted line indicates a 50% acceptance probability. The top-left insert shows the non-mimetic deep-blue model egg (right) next to natural American robin eggs (left). Numbers at the top reflect the total number of birds in each treatment.

## 4. Discussion

Host adults show elevated baseline and stress-induced corticosterone levels in response to brood parasitism [9,10]. Until now, the consequences of this endocrine response on host antiparasitic behaviours have been unknown [8]. Here we experimentally investigated if corticosterone plays a role in mediating host defences against brood parasites. We showed that treatment with mitotane, which inhibits glucocorticoid synthesis, increases the probability of acceptance of a non-mimetic deep-blue model egg by American robins, compared to a sham treatment. This is the first study to establish an endocrine basis of a widespread and effective antiparasitic host defence [7].

Hormonal mediation of host resistance against brood parasites has important implications for the ecology and evolution of these behaviours. Specifically, because of the pleiotropic effects of glucocorticoids on host phenotype [33,34; but see 35], and the sensitivity of these hormones to various environmental and social stimuli [36,37], an endocrine basis of host behaviour necessitates that we consider these defences as part of an integrated phenotype. For example, stable individual differences in glucocorticoid levels or the expression of glucocorticoid receptors, or changes in glucocorticoids in response to physical or social challenges, may explain variation in the propensity (or ability) of hosts to respond to brood parasitism. So far, the only study that has investigated such links has found a weak positive association between foreign egg acceptance and baseline corticosterone levels [25]. It is therefore unclear if the increased acceptance of model eggs in response to mitotane in this study is mediated by changes in baseline (as in [10]) or parasitism-related stress-induced [9] corticosterone levels. Indeed, our treatment validation design only allowed us to test if mitotane could alleviate a major increase in corticosterone levels in response to captivity stress. This topic therefore merits future attention.

The mechanisms through which glucocorticoids affect the probability of egg rejection are unknown. For example, a rise in glucocorticoid levels in response to a parasitic egg (or an adult brood parasite) may activate an action pattern of discrimination, recognition, and rejection [6]. Another possibility is that the effect of glucocorticoids on egg rejection is mediated by a general suppression of affiliative maternal behaviour [8]. Corticosterone has been shown to have a suppressive effect on the parental behaviour across different bird species [16,38; but see 39]. A testable hypothesis is that, in brood parasite hosts, glucocorticoids may suppress affiliative maternal behaviour towards egg-like stimuli, leading to a lower stimulus threshold for foreign-egg rejection. In contrast to this prediction, we found that nest abandonment did not depend on our treatments; however, we consider the nest abandonment to be a response to capture and injection per se, and not to the treatment type. Future studies should consider using a less capture-sensitive host species or less invasive and symmetrical manipulations (both increased vs. suppressed) of hormone levels in wild birds [40,41].

Finally, our study suggests a possible evolutionary scenario for the evolution of host defences. Hosts may first evolve endocrine-regulated tolerance or avoidance mechanisms [9,10], either in response to direct or indirect cues and costs of brood parasitism. These mechanisms may then be co-opted to regulate host defences, such as egg rejection.

In summary, we show that a widespread host defence in response to brood parasitism is mediated by glucocorticoid hormones. Future experiments should focus on the physiological and cognitive pathways that regulate this effect, e.g., testing if corticosterone mediates egg rejection through suppressing maternal behaviour in general or by activating specific cognitive action patterns. Furthermore, our research puts host defences in an integrative organismal context, therefore they should be considered in concert with the internal milieu and the stress-ecology of the host species. Finally, a natural next step is to investigate the mechanisms that underlie variation in rejection of natural parasitic eggs in hosts of egg-mimetic brood parasites, to understand how the control of host behaviour evolves in response to increasing parasitic trickery.

### Ethics

This study was approved by the animal ethics committee (IACUC) of the University of Illinois (#17259), and by permits from USA federal (MB08861A-3) and Illinois State agencies (NH19.6279). Despite our best efforts to minimize disturbance at nests, the capture and injection treatments resulted in an unanticipatedly high rate of nest abandonment (39%). This was much higher than the abandonment due to capture (but not injection) we observed in a previous season (25%) [25]. To limit negative consequences to the population, we therefore narrowed our study to a single treatment vs. the sham manipulation. Importantly, injected birds were often observed re-nesting, indicating that the effect of injections was not permanent.

## Data accessibility

The data included in these analyses will be deposited into Dryad upon acceptance.

## Authors’ contributions

MA-A and MEH designed the study, MA-A conducted field and lab work and analysed the data. MA-A and MEH co-wrote the manuscript.

## Competing interests

We declare that we have no competing interests.

## Funding

Harley Jones Van Cleave Professorship of the University of Illinois.

## Acknowledgements

We thank the landowners for permission to work on their properties and C. Goethe for assistance with field work.

